# Probabilistic forecasting in infectious disease epidemiology: The thirteenth Armitage lecture

**DOI:** 10.1101/104000

**Authors:** Leonhard Held, Sebastian Meyer, Johannes Bracher

**Author notes:** Correspondence to: Leonhard Held, Epidemiology, Biostatistics and Prevention Institute, University of Zurich, Hirschengraben 84, 8001 Zurich, Switzerland.

## Abstract

Routine surveillance of notifiable infectious diseases gives rise to daily or weekly counts of reported cases stratified by region and age group. From a public health perspective, forecasts of infectious disease spread are of central importance. We argue that such forecasts need to properly incorporate the attached uncertainty, so should be probabilistic in nature. However, forecasts also need to take into account temporal dependencies inherent to communicable diseases, spatial dynamics through human travel, and social contact patterns between age groups. We describe a multivariate time series model for weekly surveillance counts on norovirus gastroenteritis from the 12 city districts of Berlin, in six age groups, from week 2011/27 to week 2015/26. The following year (2015/27 to 2016/26) is used to assess the quality of the predictions. Probabilistic forecasts of the total number of cases can be derived through Monte Carlo simulation, but first and second moments are also available analytically. Final size forecasts as well as multivariate forecasts of the total number of cases by age group, by district, and by week are compared across different models of varying complexity. This leads to a more general discussion of issues regarding modelling, prediction and evaluation of public health surveillance data.

## 1. Introduction

Forecasting is one of the key goals of infectious disease epidemiology, and mathematical and statistical models play a central role in this task. For example, Keeling and Rohani [1] (Section 1.5) write that “models have two distinct roles, prediction and understanding. Prediction is the most obvious use of models.” However, most of traditional infectious disease epidemiology is concerned with “understanding”, and less so with forecasting.

Indeed, the World Health Organization (WHO) concluded in 2014 that “forecasting disease outbreaks is still in its infancy, however, unlike weather forecasting, where substantial progress has been made in recent years.” [2]. As a consequence, WHO has recently organized an informal consultation with more than 130 global experts entitled “Anticipating Emerging Infectious Disease Epidemics” in order “to define the elements within which epidemics of the future will occur” [3]. Likewise, the United States Federal Government has recently sponsored two prediction competitions on dengue fever in Puerto Rico and influenza-like illness in the United States^†^.

There have been various attempts to predict future infectious disease trends with mathematical or statistical models. For example, there has been success in reproducing the spread of the 2002/2003 SARS epidemic based on global flight travel data [4]. Another commonly considered problem is the prediction of influenza epidemics [5, 6, 7, 8, 9, 10]. However, these approaches often concentrate on data available as univariate time series. This is often of limited value since infectious diseases may spread differently in subgroups of the population considered, for example in different age groups. Strong spatio-temporal dynamics are also very common. A multivariate view is therefore required to predict incidence in different regions and age groups and this is the scenario we are considering in this article. Of course, other stratification levels such as gender may also be relevant in other applications.

The key requirements for stratified forecasting of infectious disease incidence are threefold: First, suitable multivariate models have to be developed for stratified time series of surveillance counts. This includes spatio-temporal models [11, 12, 13, 14] and models that reflect contact patterns between age groups [15, 16]. Secondly, focus should be on probabilistic forecasts, rather than on deterministic point predictions [17]. Probabilistic forecasting is the standard in many other areas of science, including economics and weather forecasting. The inherent difficulty in forecasting epidemics [18] makes the use of probabilistic forecasts in infectious disease epidemiology even more necessary to properly reflect forecasting uncertainty. The natural way to obtain probabilistic predictions is through the use of statistical models. An important aspect of the focus on predictions is the emphasis on the primacy of observables and the notion of a model as a (probabilistic) prediction device for such observables. Our models may therefore not perfectly represent (individual-based) disease transmission, but may still be useful for prediction of suitably aggregated surveillance data.

Thirdly, the quality of the forecasts has to be assessed with appropriate predictive model criteria. As in weather forecasting, we use proper scoring rules to assess the quality of probabilistic (count) predictions. Propriety ensures that both calibration and sharpness are addressed. Calibration is defined as the statistical consistency of the probabilistic forecasts and the observations. Sharpness refers to the concentration of the predictive distributions. Thus, the goal is to “maximize sharpness subject to calibration” [19]. Calibration of univariate forecasts can be visually assessed with probability integral transform (PIT) histograms for count data [20]. Calibration tests for count data can also be employed [21, 22, 23]. Related methods have been proposed to assess the quality of multivariate probabilistic forecasts [24, 25, 26].

With this paper we aim to review and address the above issues. In our application we focus on norovirus gastroenteritis (described in Section 2), an endemic infectious disease where the amount of historical data makes it possible to predict future disease spread. Although we apply our methods for predictive validation in infectious disease epidemiology exclusively in the `hhh4` [27] modelling framework (described in Section 3) using the R package surveillance [28], the proposed methodology for assessing probabilistic forecasts (described in Section 4) can also be applied to other modelling approaches. Section 5 presents results for the norovirus surveillance data and Section 6 concludes with some discussion.

## 2. Norovirus gastroenteritis surveillance data

Norovirus gastroenteritis is characterized by a “sudden onset of vomiting, diarrhea and abdominal cramps lasting 2–3 days” [29]. Its generation time is similar to seasonal influenza (3–4 days). The disease is highly contagious, transmitted directly from person to person, but also indirectly via contaminated surfaces or food. No vaccination is available. Weekly (lab-confirmed) counts have been downloaded from https://survstat.rki.de (Figure 1). Our training data – identical to the one analysed in Meyer and Held (2016) [16] – are stratified into 6 commonly used age groups (0–4, 5–14, 15–24, 25–44, 45–64, and 65+ years) and 12 city districts (of Berlin), and covers the period from week 2011/27 to week 2015/26. The following year (2015/27 to 2016/26) has been used to assess model predictions (test data).

**Figure 1.**
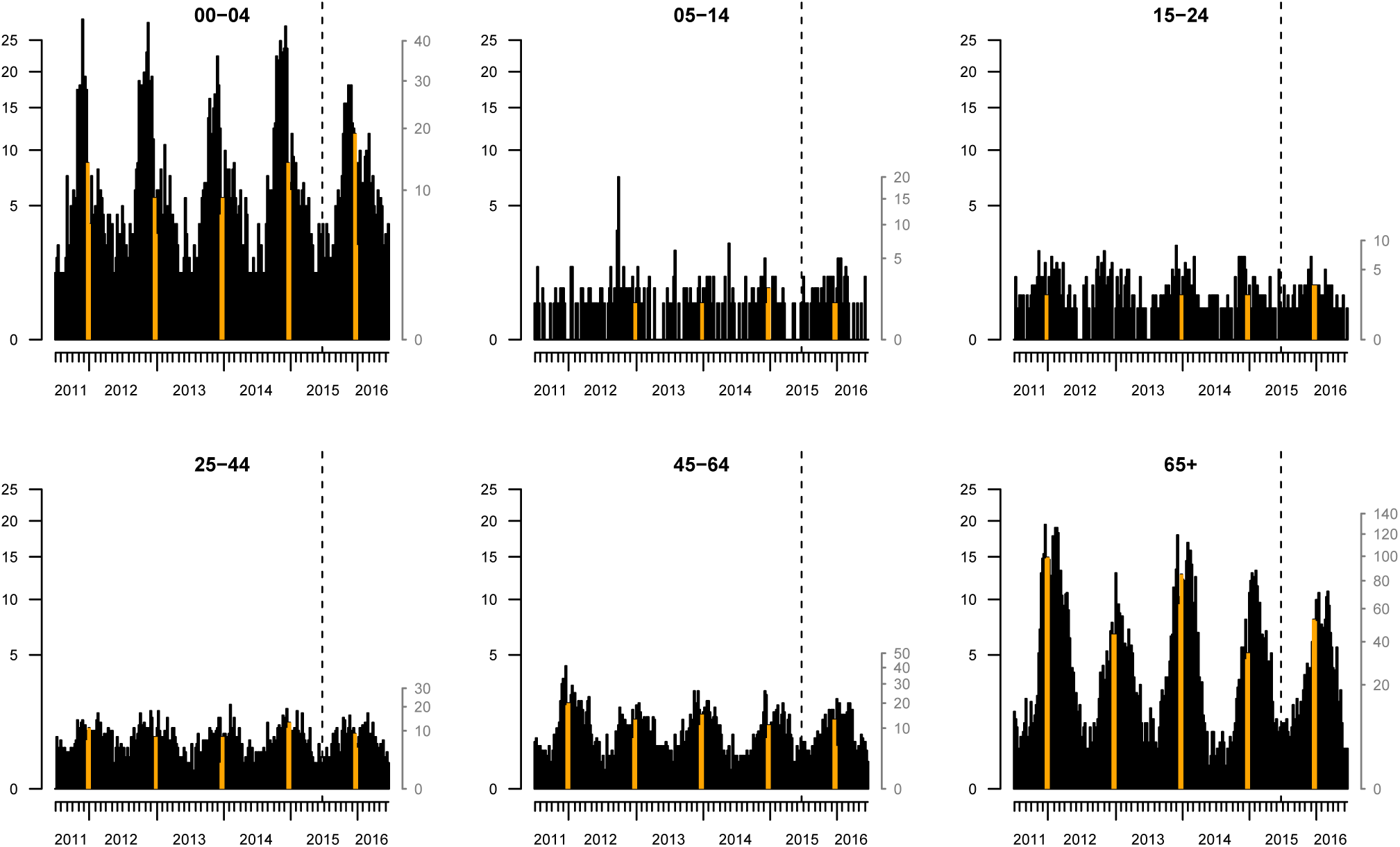
Reported incidence of norovirus infections in Berlin, Germany, stratified by age group. The incidence on the left axis is per 100 000 inhabitants and obeys the same √-scale in all panels. The corresponding counts can be read off from the right axis. The population fractions of the age groups are 4.6%, 7.8%, 10.4%, 30.4%, 27.9%, and 18.9%, respectively. The yearly Christmas break in calendar weeks 52 and 1 is highlighted. The last season (to the right of the dashed vertical line) is used as test data.

## 3. Endemic-epidemic models for infectious disease counts

We start with a modelling framework for spatio-temporal infectious disease counts in Section 3.1. This is extended in Section 3.2 to include social contact patterns between age groups.

### 3.1. Space-time model

In a series of papers [27, 30, 31, 12], a modelling framework for multivariate surveillance count time series of infectious diseases has been proposed. The formulation is built upon an additive decomposition of disease incidence into an endemic and an epidemic component. The endemic component may represent seasonal and climatic variation, heterogeneity in population numbers and other socio-demographic characteristics. The epidemic component describes the force of previously infected individuals and thus spatio-temporal interaction [12].

Let *Y*_*rt*_ denote disease counts in region *r* and week *t*. Given the counts *Y*_*r,t*−1_, *r* = 1,…,*R*, in the previous week, *Y*_*rt*_ is assumed to be negative binomial distributed, *i. e*. *Y*_*rt*_ ~ NBin(*µ*_*rt*_, *ψ*), with mean

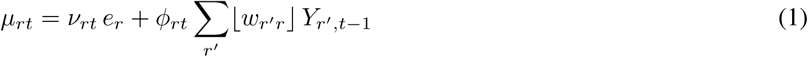

and overdispersion parameter *ψ*, so Var(*Y*_*rt*_) = *µ*_*rt*_(1 + *µ*_*rt*_). Here *ν*_*rt*_ is an unknown endemic log-linear predictor and *e*_*r*_ are known population fractions of the different districts. The epidemic log-linear predictor *ϕ*_*rt*_ describes the force of infection from time *t* − 1 to time *t* where ⌊*w*_*r′r*_⌋ denote normalized weights for *r′* to *r* transmission from *Y*_*r′,t*−1_ to *Y*_*rt*_, *i. e*. ∑_*r*_⌊*w*_*r′r*_⌋ = 1 [12].

Spatio-temporal modelling has always been an important feature of infectious disease epidemiology. For example, Keeling and Rohani [1] dedicate a whole section to spatial dispersal of infections. An important discovery was that short-time travel behaviour follows approximately a power law with respect to distance [32]. Specifically, the relative frequency Pr(*d*) of the distance *d* traversed by ~500 000 dollar bills within 4 days in the U.S. has been shown to follow a power law: Pr(*d*) ∝ d^−1.59^. An interesting feature of the power law is that it is scale-free, *i. e*. the power parameter (here −1.59) does not depend on the unit in which the distances *d* are measured. A power law for areal data has also been proposed [12, 16] and is based on the adjacency order *o*_*r′r*_ between regions *r′* and *r* as distance measure:

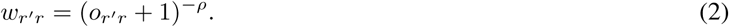

Here adjacent regions *r* and *r′* have order *o*_*r′r*_ = 1, regions where we need to traverse one other region are of order 2 and higher orders are defined accordingly. The power parameter *ρ* is treated as unknown and estimated from the data.

In our application we model *ν*_*rt*_ and *ϕ*_*rt*_ in ( 1) as

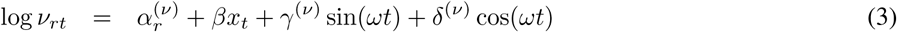

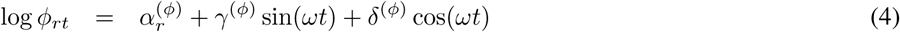

with region-specific effects 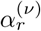 and 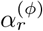 in both components, a Christmas break indicator *x*_*t*_ (to account for reduced reporting and school closure in calendar weeks 52 and 1) in the endemic component, and sinusoidal log-rates with frequency *ω* = 2π/52 in both components [31]. An alternative model replaces 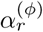 by *α*^(*ϕ*)^ + *τ* log(*e*_*r*_), so includes a parametric “gravity model” [33, 11] as described in the next section in more detail.

A multivariate branching process formulation [27] is useful to derive the (time-dependent) epidemic proportion *λ*_*t*_ as a function of the model parameters. We omit details here but will report the average epidemic proportion 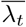, which describes the average proportion of the incidence that can be explained with the epidemic component.

### 3.2. Age-structured spatio-temporal model

Consider now counts *Y*_*grt*_ stratified by age group *g*, region *r* and week *t*. Again we use a negative binomial likelihood with mean

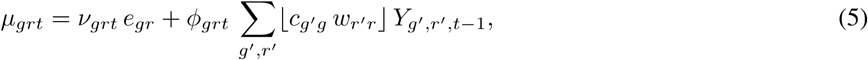

now with age group-specific overdispersion parameter *ψ*_*g*_. The unknown log-linear predictors *ν*_*grt*_ and *ϕ*_*grt*_, and the known population fractions *e*_*gr*_ also become age group-specific. There are now two sets of weights: *c*_*g′g*_ quantifies transmission from age group *g′* to age group *g*, while *w*_*r′r*_ remain spatial weights for *r′* to *r* transmission. The product *c*_*g′g*_ *w*_*r′r*_ is normalized such that ∑_*g,r*_⌊*c*_*g′g*_ *w*_*r′r*_⌋ = 1.

We now specify the model as

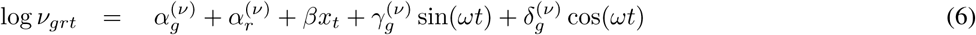

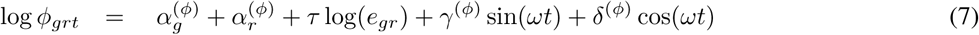

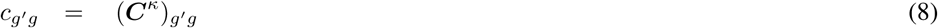

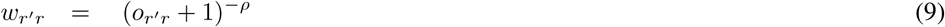

with age group and region-specific effects *α*_*g*_ and *α*_*r*_ in both components (*ν* and *ϕ*). Again we include an indicator *x*_*t*_ for the Christmas break and let seasonal terms enter both components, where we now assume age group-specific seasonality with coefficients 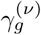 and 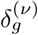 in the endemic component. Note that the formulation (7) for *ϕ*_*grt*_ extends the one used in Meyer and Held (2016) [16], since we here also allow for seasonal variation in the epidemic component [31]. The gravity model feature 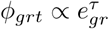 causes the amount of disease transmission to scale with the population size *e*_*gr*_ of the “importing” age group *g* in district *r* – with unknown coefficient *τ*.

To specify the weights *c*_*g′g*_, we estimate a contact matrix *C* for Germany from recorded contacts of individuals participating in the POLYMOD study [34]. Note that social contacts are reciprocal at the population level, *i. e*. there is an equal number of contacts between age group *g* and *g′* as between age group *g′* and *g*, which has been taken into account appropriately [35]. We finally specify a power transformation for the contact matrix [16],

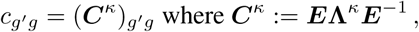

where Λ is the diagonal matrix of eigenvalues and *E* is the corresponding matrix of eigenvectors of *C*.

To investigate the relevance of the various model features, we consider four different models of increasing complexity: Model 1 is based on equation (6) with *ϕ*_*grt*_ ≡ 0 in (5), so only includes the endemic component. Model 2 extends this with an autoregression on the number of cases *Y*_*g,r,t*−1_ in the same age group and region, *i. e*. (5) reduces to *µ*_*grt*_ = *ν*_*grt*_ *e*_*gr*_ + *ϕ*_*grt*_ *Y*_*g,r,t*−1_ with *ϕ*_*grt*_ as in ( 7), but without the gravity model component *τ* log(*e*_*gr*_). Model 3 allows for spatial dispersal according to (9) and includes the gravity model in (7), but does not include contact data, so *c*_*g′g*_ = 1 for *g′* = *g*, otherwise *c*_*g′g*_ = 0. Finally, Model 4 is based on the full formulation ( 5)-( 9), so allows for dispersal across space and across age groups.

## 4. Predictive model assessment

We validate the models based on probabilistic one-step-ahead and long-term predictions. The one-step-ahead predictions are all negative binomial distributed, so the predictive mass probability function is known in closed form. Note that the underlying parameters change from prediction to prediction, as we always refit the model to all available data prior to the counts to be predicted. A simpler approach would always use the parameter estimates based on the original training data.

The model formulation implies that one-step-ahead predictions in different age groups and different regions are conditionally independent. Long-term predictions in time, sometimes called path forecasts, are generated through Monte Carlo simulation by sequentially simulating from one-step-ahead predictions. Note that our time series model is multivariate (across regions and age groups) and thus the path forecasts are also multivariate. Alternatively, the first two moments of multivariate path forecasts can be computed analytically in the model class considered, see Appendix A for details.

Predictions are computed for all combinations of 6 age groups × 12 districts × 52 weeks and have been suitably aggregated across districts, across age groups, or across time, if required. Similarly, the first two moments of aggregated counts can be calculated analytically from the moments of the original path forecasts.

### 4.1. One-step-ahead forecasts

Scoring rules (also called scoring functions) are the key measures for the evaluation of probabilistic forecasts. Scoring rules assign a numerical score based on the predictive density *f*(*y*) for the unknown quantity and on the true value *y*_*obs*_, that has later materialised. They are typically negatively oriented, *i. e*. the smaller, the better. They are called proper, if they do not provide any incentive to the forecaster to digress from her true belief, and strictly proper if any such digress results in a penalty, *i. e*. the forecaster is encouraged to quote her true belief rather than any other predictive distribution. Note that in the literature inappropriate scoring methods are still often used, *e. g*. correlation coefficients between point predictions and observations [8, 10]. It is well known in the medical literature that high correlation does not necessarily imply good agreement [36] and therefore a very poor forecasting method may have high correlation. Furthermore, usage of metrics incorporating the whole probabilistic forecast (rather than using only point predictions) is still rare [6].

For ease of presentation we drop the indices *g*, *r* and *t* in this section and denote by *Y* the predictive distribution which we compare with the actual observation *y*_*obs*_. The logarithmic score [37] is strictly proper and defined as

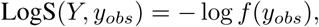

*i. e*. minus the log predictive density at the observed value. A normal approximation gives the Dawid-Sebastiani score [38]. A strictly proper alternative is the ranked probability score [19], which can be written for count data as

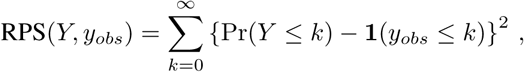

the sum of the Brier scores for binary predictions at all possible thresholds *k* ∈ {0,1,…} [20]. An equivalent definition is

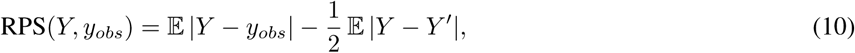

here *Y* and *Y′* are independent realisations from *f*(*y*) [19].

For Poisson and negative binomial predictive distributions, both scores (LogS and RPS) can be computed directly in the R package surveillance [28]. In order to develop a calibration test for one-step-ahead count forecasts [23], we compute the average score 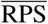 across all regions, age groups and one-step-ahead predictions and compare it to its expected value 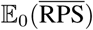 under the null hypothesis of “forecast validity” [22], sometimes also called “perfect calibration”. The difference 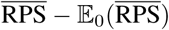 is standardized using the variance 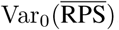. Note that our model implies that the one-step-ahead predictions in different age groups and different districts are (conditionally) independent, so under *H*_0_ independence holds for all components of 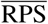 for fixed *t*. Furthermore, proofs of independence are available for the sequence of scores across time [21, 22], so in the computation of 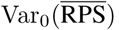 we can simply assume independence of all components of 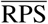. Finally, a *z*-statistic

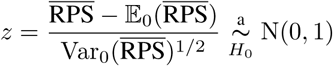

can be computed where the sign of *z* indicates if the observations are over-/underdispersed relative to the predictions (+/− sign of *z*) [23, 39]. A (two-sided) *P*-value can be computed to quantify the evidence for miscalibration.

The probability integral transform (PIT) *F*(*y*_*obs*_) is often used to visually assess calibration, here *F*(*y*) denotes the predictive CDF. Under *H*_0_, the probability integral transforms of a sequence of absolutely continuous one-step-ahead forecasts are independent and uniformly distributed [40]. Here we use a modification of the PIT histogram for count data [20]. We can also apply the Diebold-Mariano Test for equal predictive performance [41].

For negative binomial predictions, sharpness can be quantified with the estimated overdispersion parameter *ψ*, which may be age-dependent. Easier to interpret is the variance-to-mean ratio (VMR) 1 + *ψµ*, but this requires an estimate of the predictive mean *µ*, where we simply average (in each age group for age-dependent overdispersion parameters) the predictive means across weeks and districts.

### 4.2. Multivariate path forecasts

Our target quantity to assess the long-term forecasts are the number of reported cases during the test period, suitably aggregated to stratification levels of interest. In the simplest case we aggregate over week, age group and region and predict the final size, *i. e*. the total number of reported cases during the whole year. We also predict the final size in different age groups and the final size in different regions. Finally, we predict the so-called epidemic curve, *i. e*. the total number of cases for each of the 52 weeks of the test data (aggregated over age group and region), given the training data.

Predictions of aggregated counts enable us to compare our models to simpler models that are applied to suitably aggregated data. For example, we could just fit a univariate negative binomial time series model to the counts *Y*_*t*_ aggregated over age group and district to predict the total number of cases per week and to obtain final size predictions (model 5):

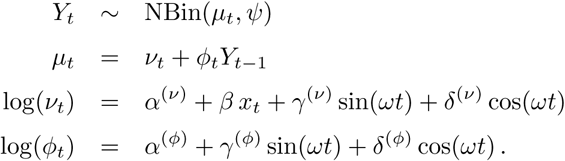

Likewise, aggregating the data over regions yields a model for *Y*_*gt*_ ~ NBin(*µ*_*gt*_, *ψ*_*g*_) to additionally obtain final size predictions in the different age groups (model 6):

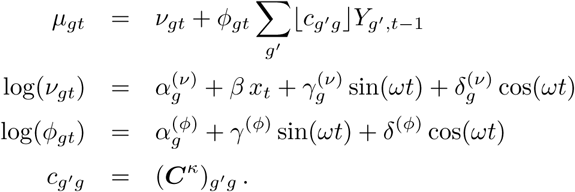

Finally, two models for regional counts *Y*_*rt*_ ~ NBin(*µ*_*rt*_, *ψ*) (aggregated over age group) have been applied. Both are based on the decomposition ( 1) with power law ( 2) and endemic model ( 3). Model 7 uses the epidemic model ( 4) with region-specific intercept 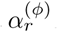. Model 8 is more parsimonious and replaces 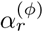 by *α*^(*ϕ*)^ + *τ* log(*e*_*r*_), so includes a gravity model component instead. Note that the more general formulation 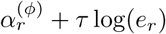 is not identifiable.

One possibility for a proper scoring rule is the multivariate Dawid-Sebastiani score [38]

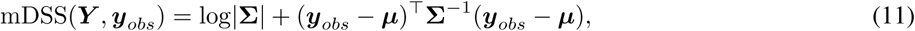

that depends only on the mean vector *µ* and the covariance matrix ∑ of the predictive distribution. The first term in ( 11) involves the determinant |∑| of the covariance matrix ∑, here ∑ is a *d* × *d* matrix. Transformed to

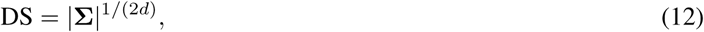

this is known as the *determinant sharpness* (DS) and recommended as a multivariate measure of sharpness [24], with smaller values corresponding to sharper predictions. In higher dimensions it is useful to report the log determinant sharpness log DS = log|∑|(2*d*) to avoid unnecessarily large numbers. For presentational reasons we also report the Dawid-Sebastiani score ( 11) in a scaled version and divide it by 2*d*. Finally, we compute an approximate *P*-value for forecast validity based on the mDSS [26]. We note that this method assumes approximate normality of the observed data, which may be questionable if the observed counts are very low.

However, evaluation of ( 11) and ( 12) based on Monte Carlo estimates of the first two moments is not recommended, since the determinant |∑| is known to be very sensitive to Monte Carlo sampling error [25]. Fortunately we can evaluate this score exactly since we have analytical results for recursive computation of *µ* and ∑ in our modelling framework, see Appendix A for details.

In order to take the full predictive distribution into account, the multivariate log-score LogS(*Y*, *y*_*obs*_) = *−*log *f*(*y*_*obs*_) will be very difficult to compute based on samples from the predictive distribution *f*(*y*). Specifically, if there is no sample exactly equal to the observed vector *y*_*obs*_, the empirical estimate of *f*(*y*_*obs*_) will be zero and the log-score will be infinite. Instead we use a generalization of the ranked probability score ( 10), the so-called energy score [19]

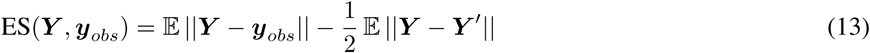

for multivariate forecasts *Y*, here ‖.‖ denotes the Euclidean norm. The energy score reduces to the Euclidean error ‖*µ* − *y*_*obs*_‖ for a deterministic point mass forecasts at *µ* and can be reported in the same unit as the observations. It has been noted [25, 42] that the ES discriminates well between forecasts with different means or variances, but less so for forecasts with different correlation structures.

Monte Carlo estimation of ( 13) is possible if samples from the predictive distribution are available [24]. These samples can also be used to derive the distribution of ES under the null hypothesis of forecast validity: Suppose we have *n*_sim_ samples *y*^(1)^,…,*y*^(*n*_sim_)^ from the predictive distribution *f*(*y*). We can then generate *n*_sim_ samples from the distribution of ES(*Y*, *y*_*obs*_) under *H*_0_ by computing the Monte Carlo approximation of (13) based on the realisation *y*_*obs*_ = *y*^(*i*)^, *i* = 1,…,*n*_sim_ and approximate the distribution of *Y* using the remaining *n*_sim_ − 1 samples from *f*(*y*). Computation of a (one-sided) Monte Carlo *P*-value is then possible based on the energy score of the actual observation *y*_*obs*_.

## 5. Results

### 5.1. Model fitting results

A comparison of selected parameters together with AIC values is given in Table 1. AIC improves for each of the features added from model 1 to model 4, where the largest impact is due to the account for autoregression (1 → 2). Likewise, the average epidemic proportion 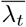 increases with increasing model complexity, so in model 4, 70% of the incidence can be explained by the epidemic component. There is strong evidence that the epidemic part scales with the population size of the “importing” districts and age groups with the coefficient *τ* estimated as 0.62 (95% CI: 0.27 to 0.97) in model 3 and 0.82 (95% CI: 0.49 to 1.14) in model 4. The power parameter *ρ* of the spatial power law ( 9) is estimated to be 2.21 (95% CI: 1.92 to 2.55) in model 3 and 2.30 (95% CI: 2.01 to 2.63) in model 4. Thus, the inclusion of social mixing between age groups in model 4 results in a slightly larger coefficient, reducing the range of spatial dispersal. Note that the results of model 4 differ slightly from the estimates reported in Meyer and Held [16] due to the additional seasonal variation in the epidemic component here. This adds two parameters but improves AIC by 18.3 points. The estimate of the power coefficient *κ* for the contact matrix *C* is 0.41 (95% CI: 0.29 to 0.60). For comparison, in model 6, the estimated exponent is nearly identical 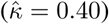 but has more uncertainty (95% CI: 0.23 to 0.67).

**Table 1.** Comparison of the different models in terms of AIC with selected parameter estimates and associated 95% Wald confidence intervals.

| Model | dim | $\Delta\text{AIC}$ | $\tau$ | $\rho$ | $\kappa$ | $\bar{\lambda}_t$ |
| --- | --- | --- | --- | --- | --- | --- |
| 1 | 36 | 0.0 | — | — | — | 0.00 |
| 2 | 55 | -514.4 | — | — | — | 0.48 |
| 3 | 57 | -631.1 | 0.62 (0.27–0.97) | 2.21 (1.92–2.55) | — | 0.68 |
| 4 | 58 | -677.7 | 0.82 (0.49–1.14) | 2.30 (2.01–2.63) | 0.41 (0.29–0.60) | 0.70 |

### 5.2. Assessing one-step-ahead forecasts

Figure 2 gives the variance-to-mean ratios of the one-step-ahead forecasts in the different age groups and models. Generally the VMR decreases with increasing model complexity. The predictions become sharper in age groups 65+ and 45–64 when we move from model 1 to 2, 3, and finally to model 4. In age group 00–04 we also see sharper predictions in model 2–4 compared to model 1.

The mean RPS with *z*-value and *P*-value is shown in Table 2. The PIT histograms [20] based on all 6 × 12 × 52 = 3744 one-step-ahead forecasts are shown in Figure 3. The mean RPS values are in the expected order with smaller (*i. e*. better) values for models of increasing complexity and the full model 4 as the best. For all models except for model 1 we see some evidence for underdispersed predictions (Figure 3) with more PITs close to one than we would expect under forecast validity. This indicates potential problems of the negative binomial distribution in the more complex model formulations.

**Figure 2.**
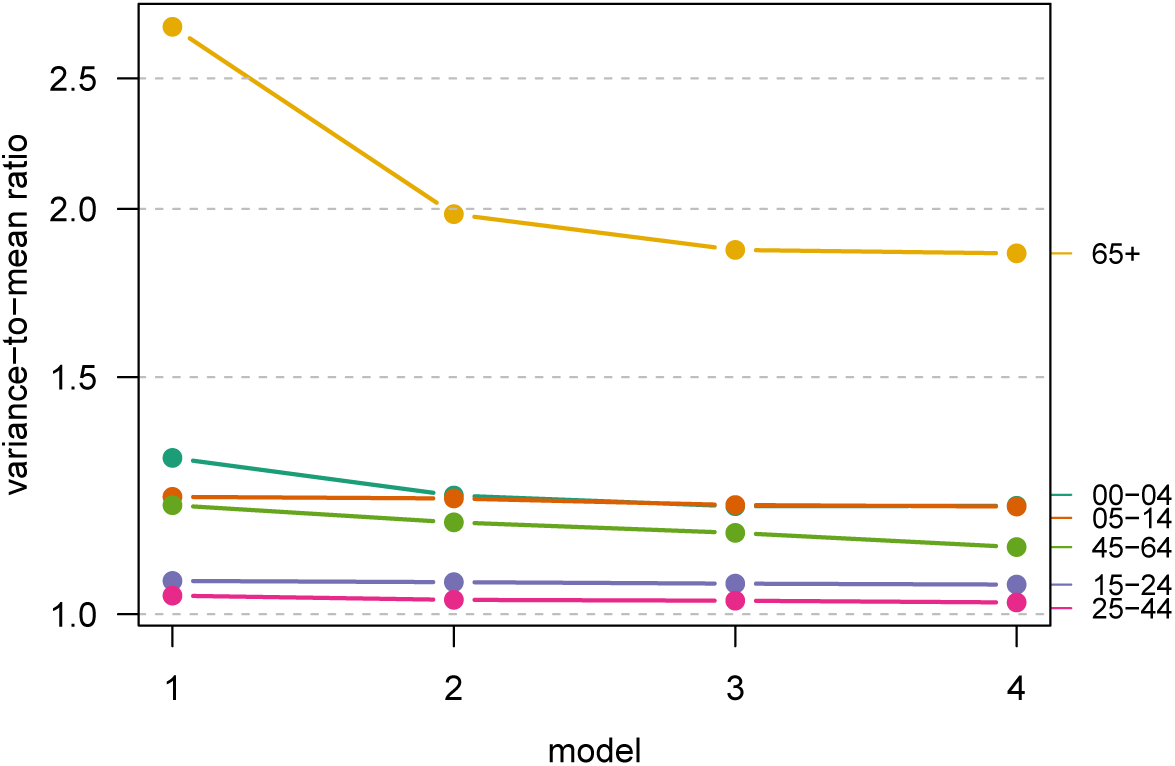
Estimated age group-specific variance-to-mean ratios of the one-step-ahead predictive distributions.

**Table 2.** Comparison of 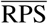 and calibration test results, overall and in the first and last age group.

| Model | Overall |  |  | Age group 00–04 |  |  | Age group 65+ |  |  |
| --- | --- | --- | --- | --- | --- | --- | --- | --- | --- |
| | $\overline{\text{RPS}}$ | $z$ | $P$-value | $\overline{\text{RPS}}$ | $z$ | $P$-value | $\overline{\text{RPS}}$ | $z$ | $P$-value |
| 1 | 0.454 | 0.65 | 0.52 | 0.535 | 2.84 | 0.005 | 1.112 | -1.83 | 0.067 |
| 2 | 0.436 | 1.35 | 0.18 | 0.516 | 2.39 | 0.017 | 1.019 | -0.17 | 0.87 |
| 3 | 0.434 | 1.80 | 0.072 | 0.513 | 2.18 | 0.029 | 1.009 | 0.82 | 0.41 |
| 4 | 0.433 | 2.28 | 0.022 | 0.512 | 2.26 | 0.024 | 1.008 | 0.79 | 0.43 |

The corresponding results for age groups 00–04 and 65+ are also shown in Table 2. In age group 00–04, there is evidence for underdispersed predictions with values of the *z*-statistic larger than 2 for all four models and an even more pronounced peak of the PIT histogram close to one than for the overall set of forecasts (compare the middle panel in Figure 5). However, we see some evidence for overdispersed predictions in age group 65+ for model 1 with a negative value of the *z*-statistic, which fits the corresponding pattern of the PIT histogram shown in the right panel of Figure 5.

**Table 3.** *P*-values from the Diebold-Mariano Test for the pairwise comparison of 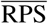 across the different models.

|  | model 1 | model 2 | model 3 | model 4 |
| --- | --- | --- | --- | --- |
| model 1 | — | < 0.0001 | < 0.0001 | < 0.0001 |
| model 2 | — | — | 0.48 | 0.23 |
| model 3 | — | — | — | 0.12 |
| model 4 | — | — | — | — |

*P*-values from the Diebold-Mariano Test are shown in Table 3. We see very strong evidence (*p* < 0.0001) for differences in predictive performance between model 1 and the other three models but no evidence for differences in predictive performance between models 2, 3 and 4.

**Figure 3.**
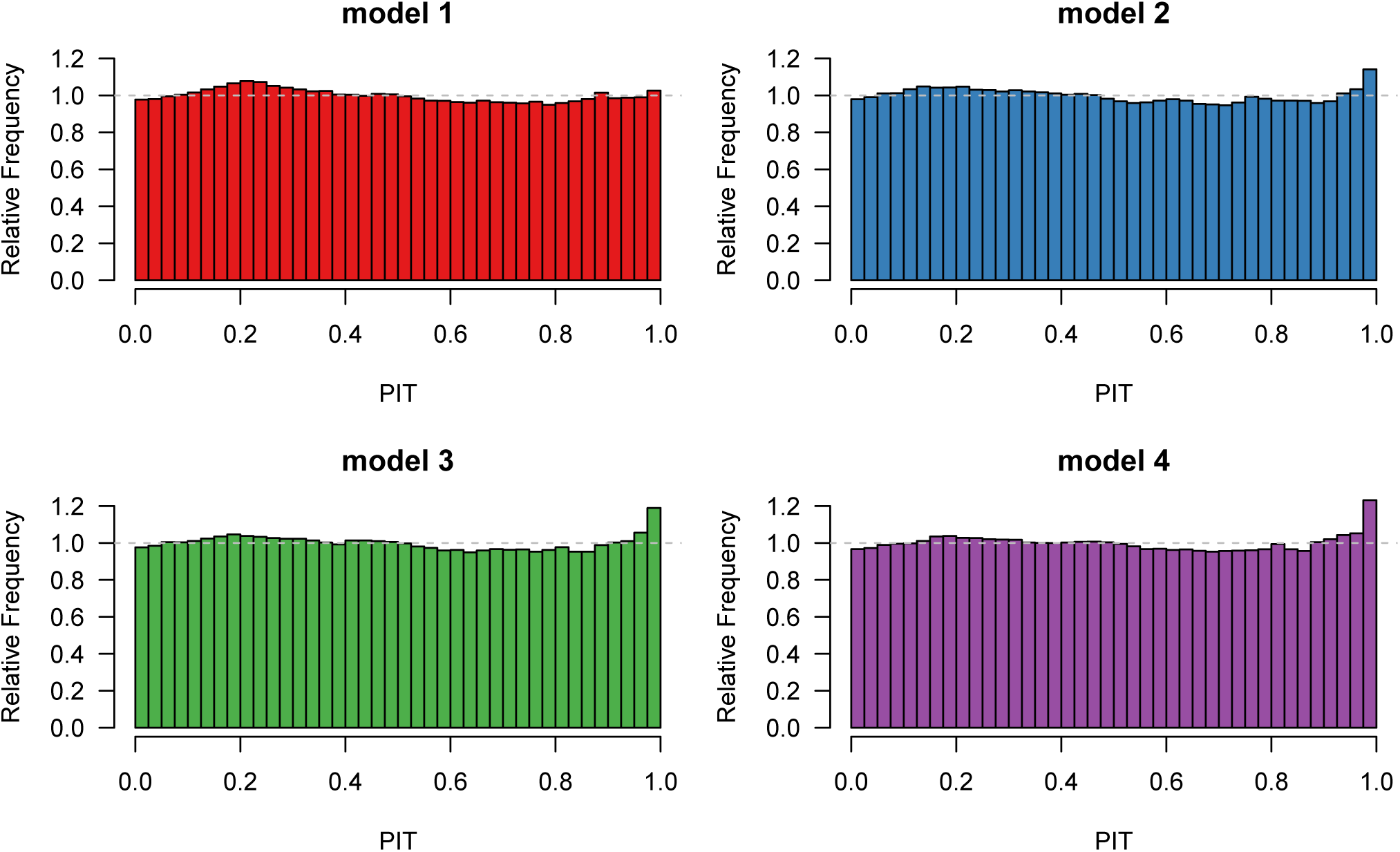
PIT histograms of the 3744 one-step-ahead forecasts for the different models.

### 5.3. Assessing long-term forecasts

Final size predictions are shown in Figure 4. All models tend to overpredict the total number of cases with varying uncertainty. In the worst cases (models 1, 2 and 8), the observed number of cases is not or only poorly supported by the predictions. The best model (in terms of RPS) is model 4, the most complex formulation, followed by model 6. This underlines the importance of using an age-structured modelling approach supported by social contact data.

Figure 5 (left panel) displays the probabilistic forecasts for the total number of cases in the different age groups for models 1–4 and 6. While all models predict the number of cases reasonably well in the lower five age groups, there are remarkable differences for the last age group 65+. Model 1 overpredicts the number of cases in age group 65+ considerably, while models 2–4 give similar point predictions, but with increasing uncertainty. As a consequence, the observed number of cases in age group 65+ is well supported by the predictive distributions of models 3, 4 and 6, less so for model 2, and not at all for model 1.

The middle and right panels of Figure 5 give PIT histograms of the one-step-ahead forecasts for the five models for age group 00–04 and 65+, respectively. The PIT histograms for the other age groups are all very close to uniformity. The PIT histograms for age group 00–04 are very similar for models 1–4 with a pronounced peak at values close to 1. This indicates a tendency to underpredict relatively large observed values. A similar pattern can be seen for model 6, where the PIT histogram is based on a smaller number of observations. The PIT histogram for age group 65+ and model 1 indicates a bias of the estimates with a tendency to predict larger values than observed. This results in overprediction of the aggregated counts shown in the left panel. In contrast, the PIT histogram of model 6 is inconspicuous. The other three models compensate the bump of the PIT histogram around 0.2 with another peak at values of PIT close to 1. This seems to lead to greater uncertainty of the final size predictions as shown in the left panel. Note that the calibration tests based on the mean RPS of the one-step-ahead forecasts (Table 2) give evidence for miscalibrated forecasts in age group 00–04 for all the models, but no such evidence for age group 65+. However, the negative value of the *z*-statistic of model 1 (with corresponding *P*-value of 0.067) already indicated a potential problem of this formulation.

**Figure 4.**
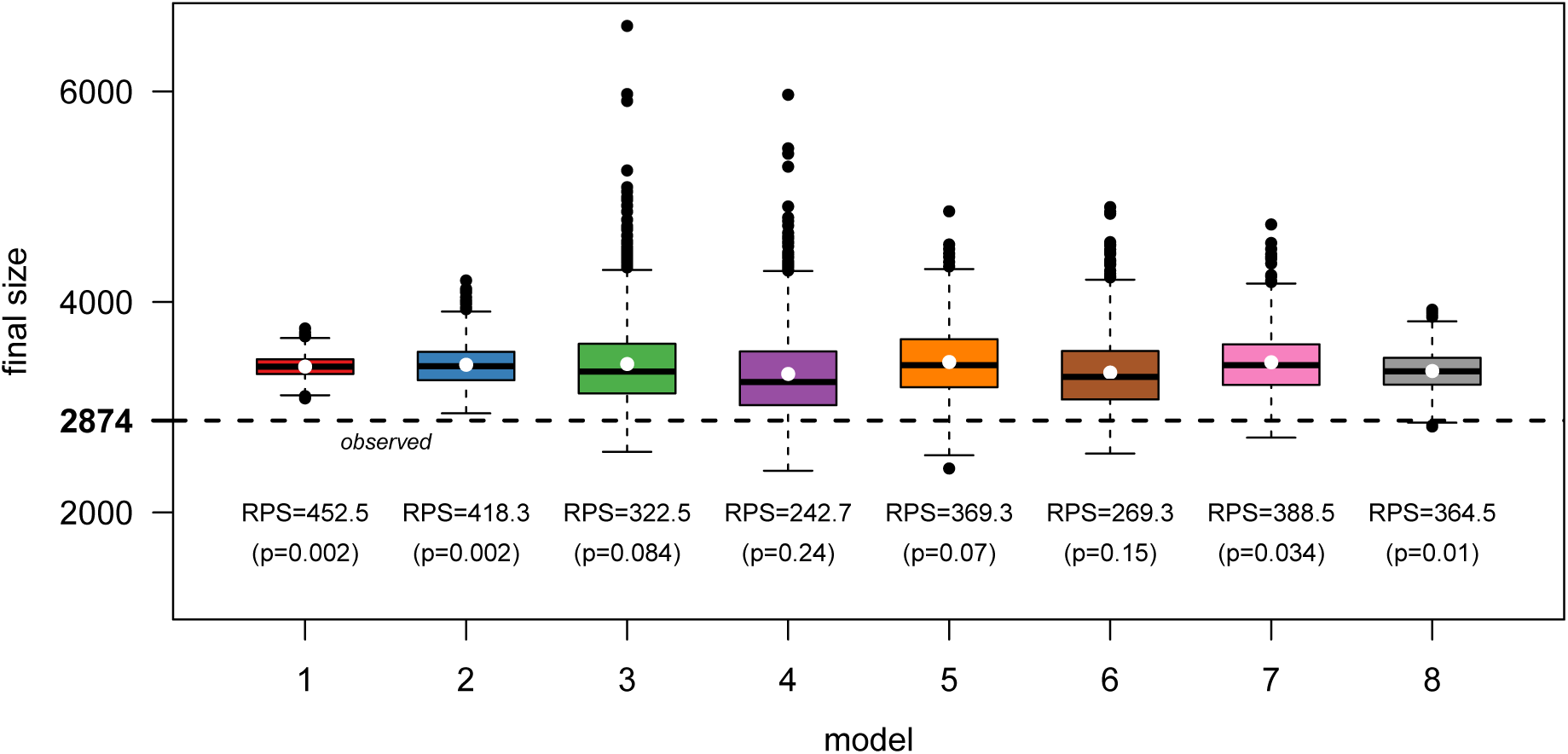
Final size predictions based on 1000 simulations with RPS and two-sided *P*-value. The white dots represent the predictive means.

Table 4 gives values of the log determinant sharpness for the different models and the different stratification levels. The first column gives values for total final size, where we observe for models 1 to 4 a tendency for larger values of logDS, *i. e*. more dispersed predictions, with increasing complexity, see also Figure 4. However, this pattern is less clear for the models 5–8 that work on aggregated data. An interesting pattern appears for the predictions by age group, region, or week: With only one exception, the models that analyse the data in the finest resolution (1–4) give sharper predictions than the models that analyse aggregate data (5–8). In particular, the most sophisticated model with both a spatial power law and social contact data (model 4) gives the sharpest prediction for the epidemic curve. This can be explained by the fact that the predictions from this model have the largest autocorrelations among all models from week 20 onwards (and also quite large before week 20), see Figure 6 bottom. Note also that the logDS values by region roughly reflect the order of the correlations between regions (Figure 6 top right) with a tendency for larger values of logDS for smaller correlations. However, logDS is also a function of the marginal variances, so the agreement of logDS and the correlations is not perfect. Similarly, the correlations of the predictions between age groups, shown in the top left panel of Figure 6 for models 4 and 6, do not lead to smaller values of logDS by age group because of larger variances compared to models 1 and 2, as can be seen in the left panel of Figure 5.

**Table 4.** Log determinant sharpness (logDS).

| Model | Total |  | by Age Group |  | by Region |  | by Week |  |
| --- | --- | --- | --- | --- | --- | --- | --- | --- |
|  | logDS | rank | logDS | rank | logDS | rank | logDS | rank |
| 1 | 4.63 | 1 | 3.12 | 1 | 3.30 | 1 | 2.30 | 4 |
| 2 | 5.32 | 3 | 3.32 | 2 | 3.78 | 6 | 2.29 | 2 |
| 3 | 6.06 | 7 | 3.59 | 4 | 3.62 | 2 | 2.29 | 3 |
| 4 | 6.10 | 8 | 3.51 | 3 | 3.63 | 3 | 2.27 | 1 |
| 5 | 5.86 | 5 |  |  |  |  | 2.57 | 8 |
| 6 | 5.92 | 6 | 3.64 | 5 |  |  | 2.41 | 7 |
| 7 | 5.68 | 4 |  |  | 3.66 | 4 | 2.32 | 6 |
| 8 | 5.21 | 2 |  |  | 3.69 | 5 | 2.30 | 5 |

**Figure 5.**
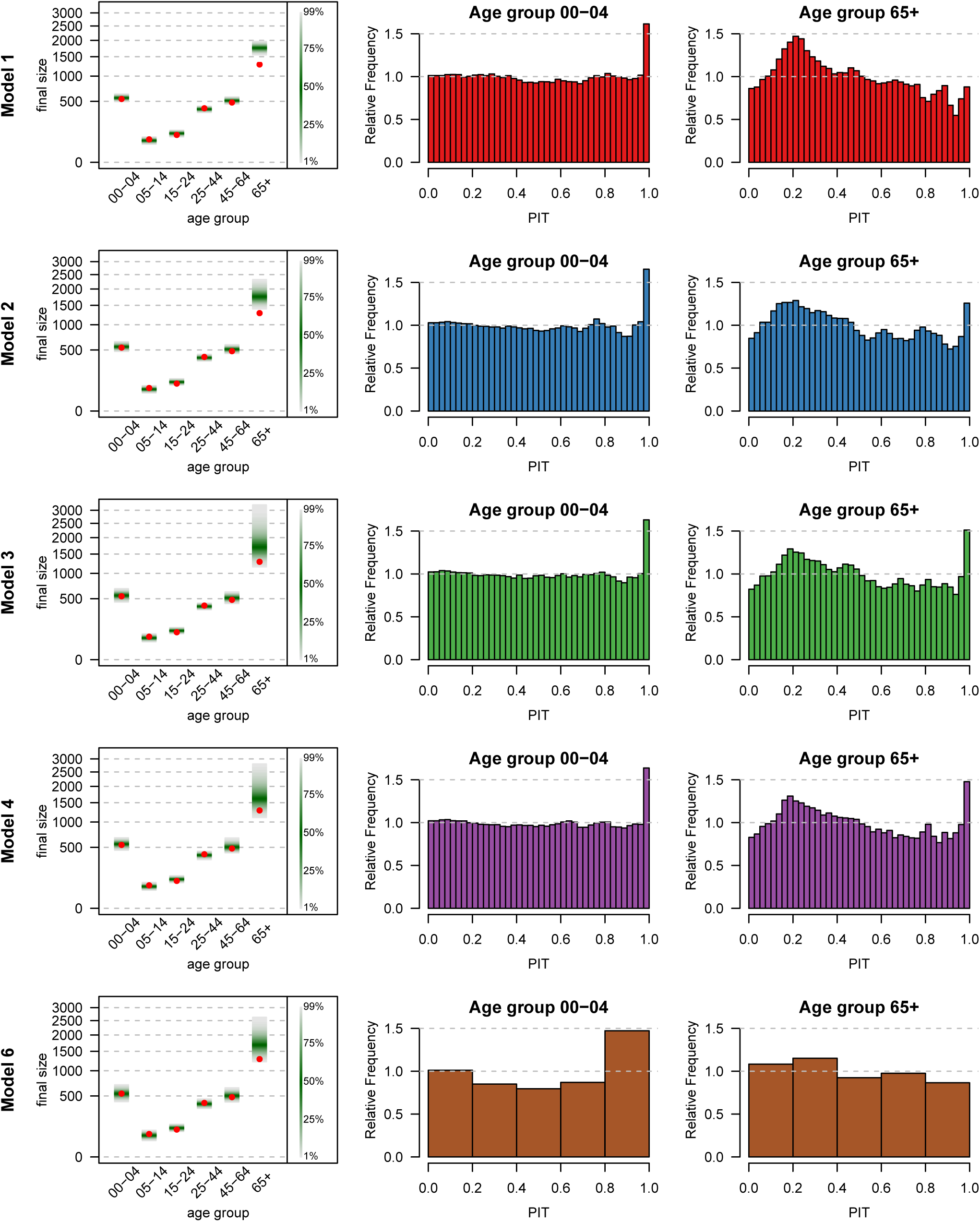
Left panel: Final size predictions in the different age groups based on 1000 simulations from models 1–4 and 6. Middle and right panel: PIT histograms of one-step-ahead forecasts in age groups 00–04 and 65+, respectively. The PIT histograms for model 6 are based on 52 observations each, whereas the other histograms are based on 12 × 52 observations.

**Figure 6.**
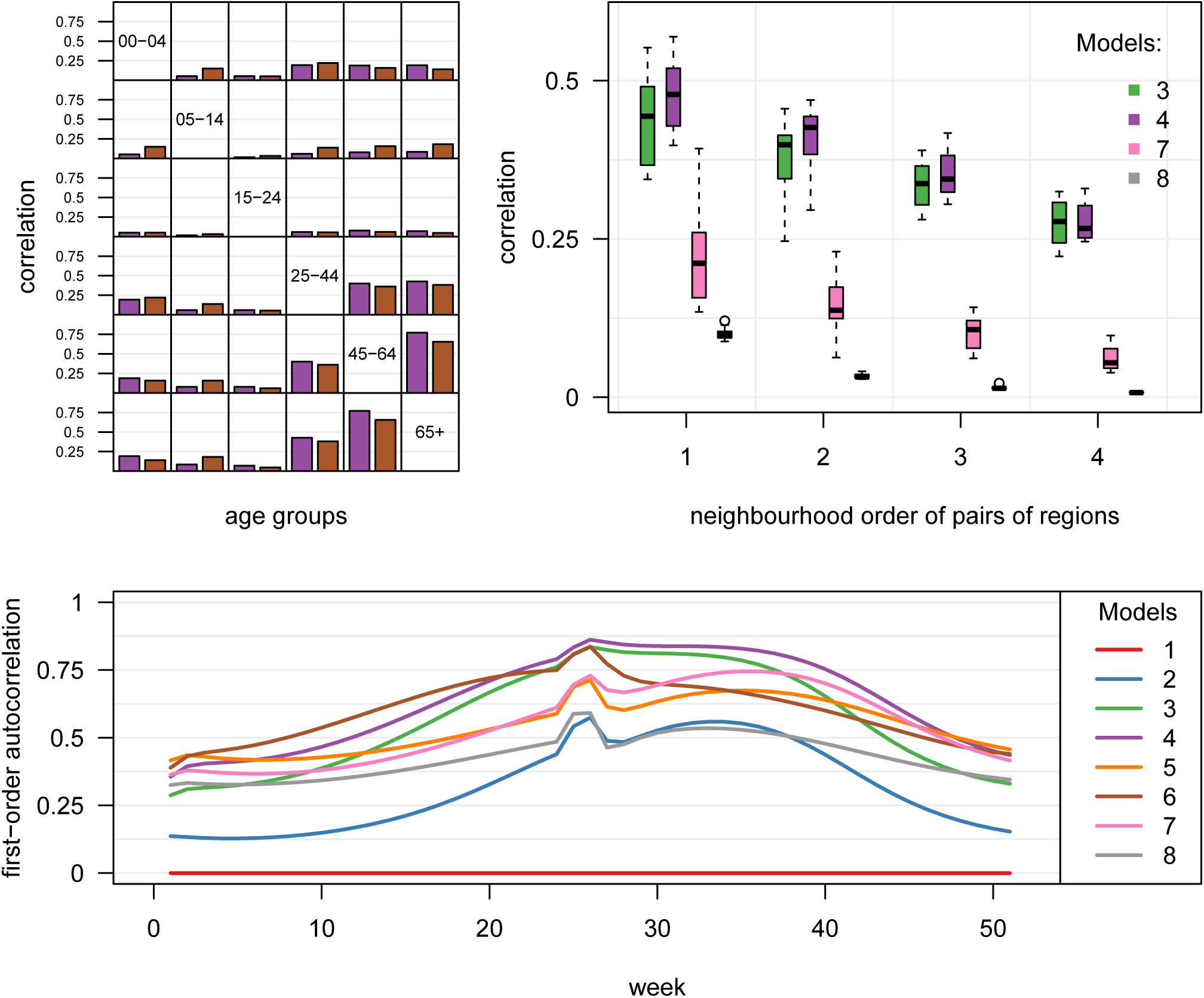
Correlations implied by the different models: between age-groups (top left), regions (top right) and consecutive weeks (bottom). Models where correlations are 0 by construction are omitted from the two top figures.

Table 5 gives both the energy and the Dawid-Sebastiani score by different stratification levels. Likewise, Table 6 gives the corresponding *P*-values for forecasts validity. The two scores agree quite well with model 4 giving the best (or second best) predictions in total, by age group and by region. Somewhat surprisingly, model 4 gives quite poor predictions by week for the DSS (rank 7), but not for the ES (rank 2). This can be explained by the fact that the ES is known to be insensitive to misspecifications in the correlation structure of multivariate predictions. We have already noted that the autocorrelations of the model 4 predictions are quite large. However, the observed time series (see Figure 7), has an oscillating pattern around weeks 20 and 30. This discrepancy between observed and predicted correlation seems to be detected by the DSS, but not by the ES. This can also be seen from Table 6, where the test based on DSS gives strong evidence for miscalibration of the forecasts by week for all models except for models 5 and 6. Interestingly, these two models have the largest values of logDS, see Table 4, but are still best in terms of DSS, see Table 5. Larger uncertainty of the predictions (see Figure 7) combined with relatively small autocorrelations (see bottom panel of Figure 6) seems to make model 5 the best in terms of DSS. Note that the corresponding test based on the ES identifies only model 1 and 8 as miscalibrated (and model 2 to a lesser extent). For the other stratification levels the *P*-values based on the DSS tend to be larger than the corresponding ones based on the ES.

**Figure 7.**
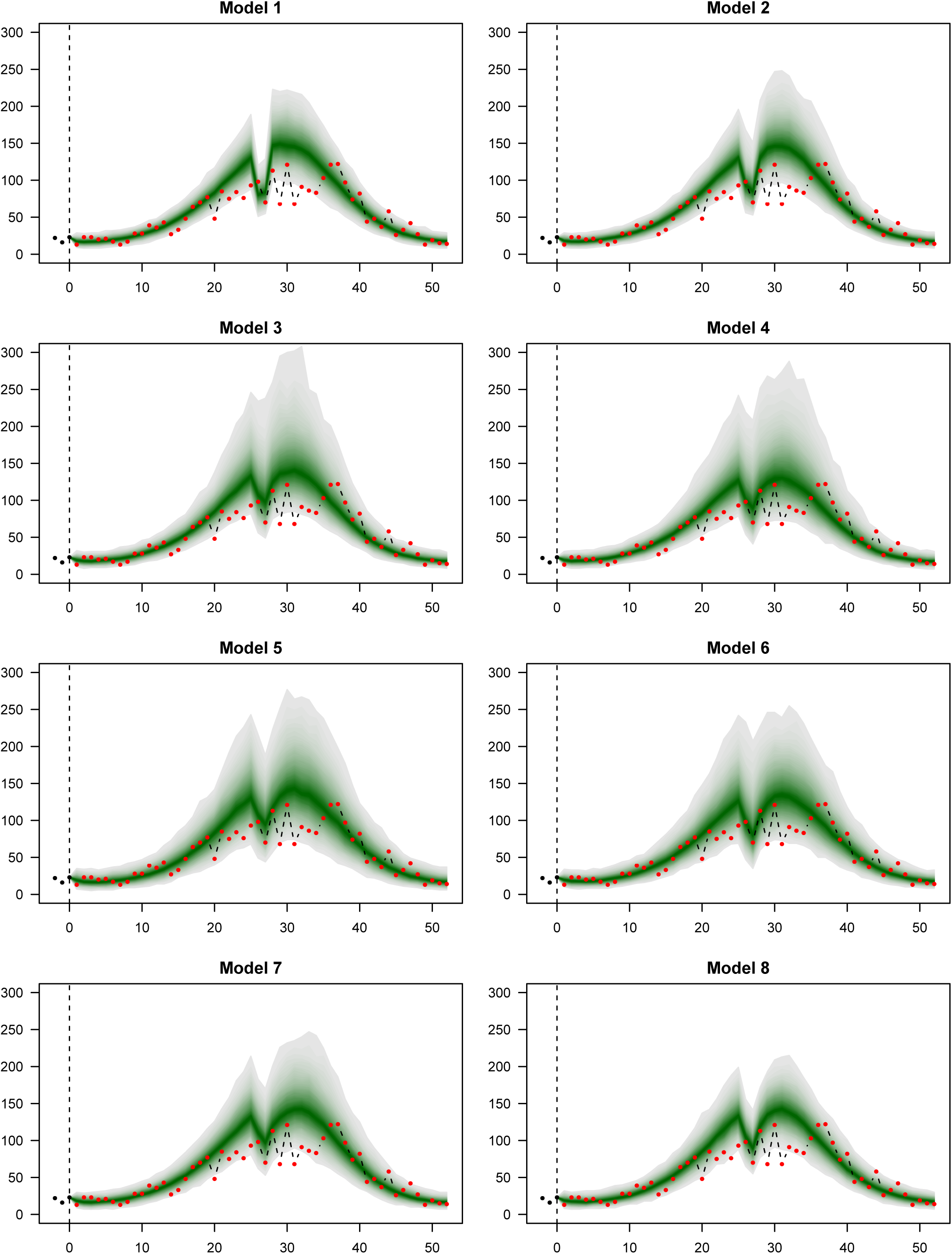
Long-term predictions of the epidemic curve in the last 52 weeks.

**Table 5.** Energy score and scaled Dawid-Sebastiani score for the different models and aggregation levels.

| Model | Total |  |  |  | by Age Group |  |  |  | by Region |  |  |  | by Week |  |  |  |
| --- | --- | --- | --- | --- | --- | --- | --- | --- | --- | --- | --- | --- | --- | --- | --- | --- |
|  | ES | rank | DSS | rank | ES | rank | DSS | rank | ES | rank | DSS | rank | ES | rank | DSS | rank |
| 1 | 452.5 | 8 | 17.10 | 8 | 409.7 | 5 | 5.67 | 5 | 161.8 | 4 | 4.85 | 6 | 136.3 | 8 | 3.32 | 8 |
| 2 | 418.3 | 7 | 8.67 | 7 | 377.3 | 4 | 4.13 | 4 | 162.7 | 5 | 4.25 | 5 | 133.5 | 7 | 3.12 | 4 |
| 3 | 322.5 | 3 | 6.83 | 3 | 293.0 | 3 | 3.93 | 2 | 158.8 | 3 | 3.98 | 1 | 126.2 | 6 | 3.14 | 6 |
| 4 | 242.7 | 1 | 6.58 | 1 | 227.0 | 1 | 3.88 | 1 | 132.8 | 1 | 3.99 | 2 | 109.3 | 2 | 3.21 | 7 |
| 5 | 369.3 | 5 | 7.03 | 4 |  |  |  |  |  |  |  |  | 118.3 | 3 | 3.05 | 1 |
| 6 | 269.3 | 2 | 6.68 | 2 | 277.3 | 2 | 4.07 | 3 |  |  |  |  | 108.0 | 1 | 3.07 | 2 |
| 7 | 388.5 | 6 | 7.39 | 5 |  |  |  |  | 180.2 | 6 | 4.10 | 3 | 122.4 | 4 | 3.13 | 5 |
| 8 | 364.5 | 4 | 8.41 | 6 |  |  |  |  | 136.1 | 2 | 4.22 | 4 | 123.5 | 5 | 3.11 | 3 |

**Table 6.** *P*-values for forecast validity.

| Model | Total |  | by Age Group |  | by Region |  | by Week |  |
| --- | --- | --- | --- | --- | --- | --- | --- | --- |
|  | Direct | DSS | ES | DSS | ES | DSS | ES | DSS |
| 1 | 0.002 | < 0.0001 | < 0.0001 | < 0.0001 | 0.0005 | 0.0002 | 0.0002 | < 0.0001 |
| 2 | 0.002 | 0.01 | 0.021 | 0.14 | 0.17 | 0.52 | 0.034 | 0.002 |
| 3 | 0.084 | 0.21 | 0.19 | 0.66 | 0.16 | 0.72 | 0.15 | 0.001 |
| 4 | 0.24 | 0.33 | 0.27 | 0.62 | 0.29 | 0.72 | 0.25 | 0.0001 |
| 5 | 0.07 | 0.13 |  |  |  |  | 0.35 | 0.57 |
| 6 | 0.15 | 0.22 | 0.14 | 0.52 |  |  | 0.34 | 0.064 |
| 7 | 0.034 | 0.064 |  |  | 0.097 | 0.57 | 0.089 | 0.003 |
| 8 | 0.01 | 0.011 |  |  | 0.093 | 0.39 | 0.008 | 0.003 |

## 6. Discussion

In this paper we have described a multivariate model framework for infectious disease surveillance counts by borrowing strength from different regions and different age groups. This model framework provides probabilistic forecasts that are useful for epidemic forecasting. Specifically, means and covariance matrices are available analytically as well as Monte Carlo samples from the full predictive distribution. Predictive model assessment helps to identify poor predictive models. The application has shown the importance of predictive model assessment in different strata. We have emphasized the importance to use appropriate methodology for predictive model assessment, including proper scoring rules and tests for forecast validity.

Our study has shown that complex modelling on the original fine resolution generally leads to better predictions of future disease incidence, even of aggregated quantities. As a consequence, the most complex model 4 was nearly always the best in predictive performance. The only exception are predictions by week, where model 4 turned out to predict poorly in terms of DSS. We were able to explain this feature by the discrepency of strong predicted correlations on the one hand, but oscillating observed number of cases on the other hand. This model deficiency was not detected by the energy score, in accordance with similar observations made in the literature [25, 42]. Consideration of models for aggregated data showed the importance to integrate contact pattern data in the analysis of age-stratified surveillance data (model 6), as this model was consistently among the top three models in terms of ES and DSS. The spatial dimension turned out to be less important, which may be different in applications with more districts.

Our forecasts were always based on one particular model. A possible extension of our approach would be to use a Bayesian model average framework to combine forecasts from different models into one averaged forecast. The model weights may even differ based on differing forecasts and could be based on AIC or BIC computed from the training data [43], if the models considered act on the same data resolution. Such model averages are known to have better forecast properties and have been successfully applied in weather forecasting [44].

## Appendix A

We derive the first two moments of multivariate path forecasts in the hhh4 modelling framework. For this purpose, it is irrelevant whether strata are defined by age groups, regions or both, so we only use one index *r* = 1,…,*R* to denote strata. The weeks are indexed such that *t* = 0 corresponds to the week on which we condition our path forecasts. Since the following derivations do not rely on model-specific choices for offsets or weights, we generalise (1) as

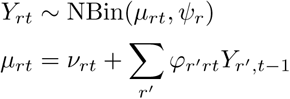

where *φ*_*r′rt*_ is a function of *ϕ*_*rt*_ and *w*_*r′r*_.

We can derive several recursive relationships for the conditional moments of 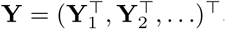, where Y_*t*_ = (*Y*_1*t*_, *Y*_2*t*_,…,*Y*_*Rt*_)^⊤^. For the means we obtain

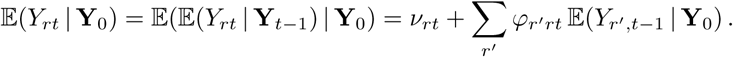

Similarly we can derive the second moments

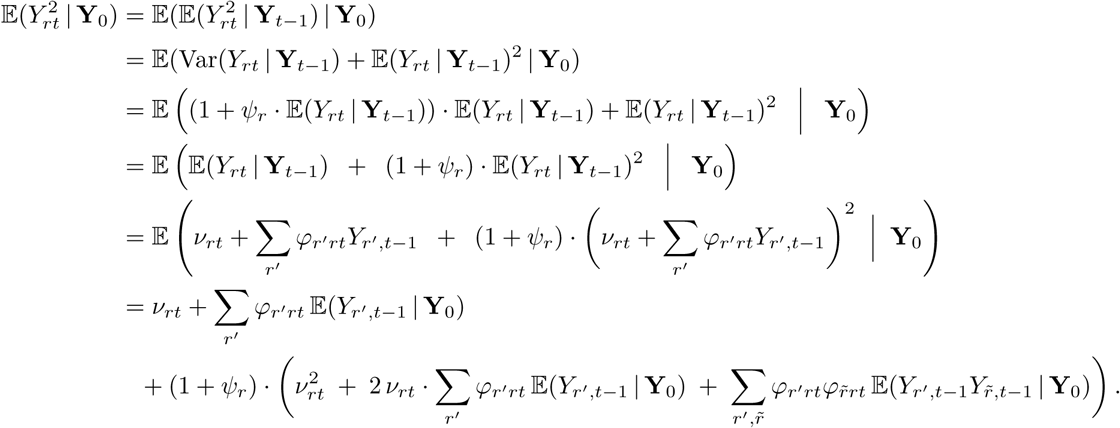

A somewhat simpler result can be obtained for the product terms (*r ≠ s*)

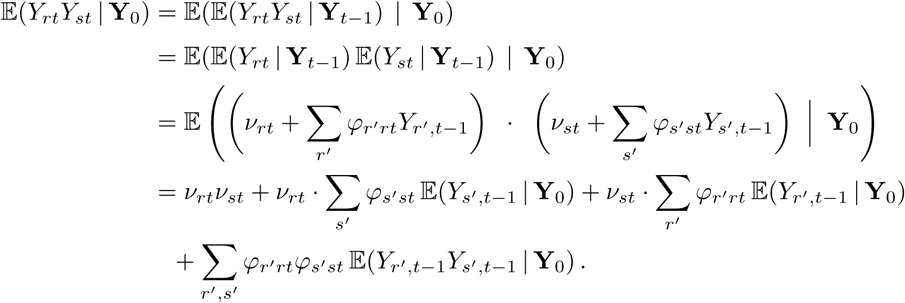

These componentwise formulations are somewhat involved, but a nice matrix formulation of the recursion exists. Define 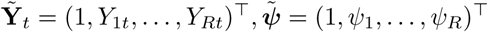 and

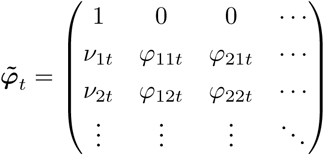

such that

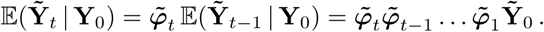

With

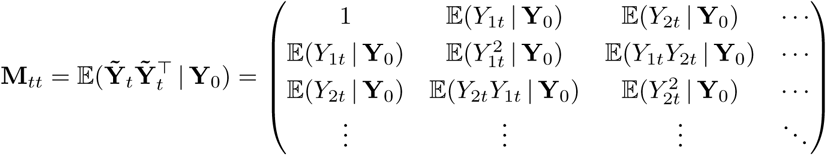

the above moment equations then allow for a simple recursive procedure consisting of one matrix transformation and one modification of the diagonal:

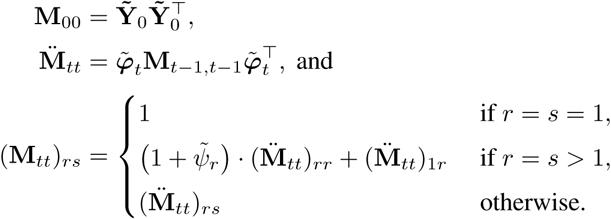

This is still limited to pairs of counts from the same week *t*. The following allows calculations across weeks:

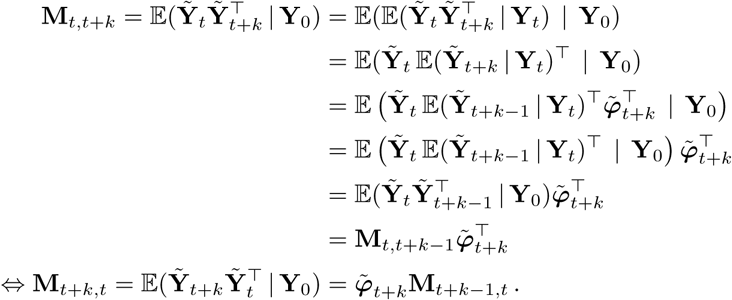

To compute 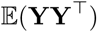 from the M_*tu*_’s we only need to omit the first row and column of each M_*tu*_. The entire recursive construction of this matrix can be visualized as follows:

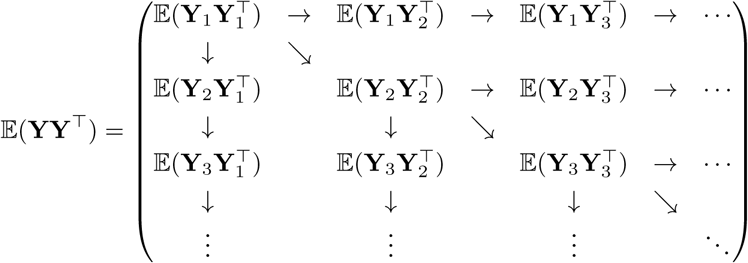

The covariance matrix is then calculated as 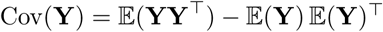.

To aggregate across strata (*e. g*. to break our results down to stratification only by region, only by age group etc.), we can use Cov(BY) = B Cov(Y)B^⊤^ where the matrix B is suitably defined.

## Acknowledgements

This paper summarizes and extends the 13th Armitage Lecture, presented by the first author at the Medical Research Council Biostatistics Unit in Cambridge, UK on November 12, 2015. A videotaped version can be found at http://www.statisticsviews.com/details/webinar/8526701/Armitage-Lecture-2015.html. The support and hospitality of the Biostatistics Unit are gratefully acknowledged. We also acknowledge helpful comments by Phil Dawid, in particular for making us aware of his work on calibration tests [21, 22]. The authors’ research was supported by the Swiss National Science Foundation (project #137919).

## Footnotes

† https://predict.phiresearchlab.org/

## References

1. Keeling MJ, Rohani P. Modeling Infectious Diseases in Humans and Animals. Princeton University Press, 2008.

2. World Health Organization. Anticipating epidemics. Weekly Epidemiological Record 2014; 89(22):244. URL http://www.who.int/wer.

3. World Health Organization ( (ed.)). Anticipating Emerging Infectious Disease Epidemics: Meeting report of WHO informal consultation, WHO Press: Geneva, Switzerland, 2016. URL http://www.who.int/csr/disease/anticipating_epidemics/meeting-report-2015/en/.

4. Hufnagel L, Brockmann D, Geisel T. Forecast and control of epidemics in a globalized world. Proceedings of the National Academy of Sciences of the United States of America 2004; 101:15 124–15 129, doi:10.1073/pnas.0308344101.

5. Birrell PJ, Ketsetzis G, Gay NJ, Cooper BS, Presanis AM, Harris RJ, Charlett A, Zhang XS, White PJ, Pebody RG, et al.. Bayesian modeling to unmask and predict influenza A/H1N1pdm dynamics in London. Proceedings of the National Academy of Sciences of the United States of America 2011; 108(45):18 238–18 243, doi:10.1073/pnas.1103002108.

6. Chretien JP, George D, Shaman J, Chitale RA, McKenzie FE. Influenza forecasting in human populations: A scoping review. PLOS ONE 2014; 9(4):e94 130, doi:10.1371/journal.pone.0094130.

7. Nsoesie E, Mararthe M, Brownstein J. Forecasting peaks of seasonal influenza epidemics. PLOS Currents Outbreaks 2013; 5, doi:10.1371/currents.outbreaks.bb1e879a23137022ea79a8c508b030bc.

8. Santillana M, Nguyen AT, Dredze M, Paul MJ, Nsoesie EO, Brownstein JS. Combining search, social media, and traditional data sources to improve influenza surveillance. PLOS Computational Biology 2015; 11(10):e1004 513, doi:10.1371/journal.pcbi.1004513.

9. Dukic V, Lopes HF, Polson NG. Tracking epidemics with Google Flu Trends data and a state-space SEIR model. Journal of the American Statistical Association 2012; 107(500):1410–1426, doi:10.1080/01621459.2012.713876.

10. Yang S, Santillana M, Kou SC. Accurate estimation of influenza epidemics using Google search data via ARGO. Proceedings of the National Academy of Sciences of the United States of America 2015; 112(47):14 473–14 478, doi:10.1073/pnas.1515373112.

11. Xia Y, Bjørnstad ON, Grenfell BT. Measles metapopulation dynamics: A gravity model for epidemiological coupling and dynamics. The American Naturalist 2004; 164(2):267–281, doi:10.1086/422341.

12. Meyer S, Held L. Power-law models for infectious disease spread. The Annals of Applied Statistics 2014; 8(3):1612–1639, doi:10.1214/14-AOAS743.

13. Riley S, Eames K, Isham V, Mollison D, Trapman P. Five challenges for spatial epidemic models. Epidemics 2015; 10:68–71, doi:10.1016/j.epidem.2014.07.001.

14. Höhle M. Infectious Disease Modelling. Handbook of Spatial Epidemiology, Lawson AB, Banerjee S, Haining RP, Ugarte MD (eds.). chap. 26, Chapman & Hall/CRC Handbooks of Modern Statistical Methods, Chapman and Hall/CRC: Boca Raton, 2016; 477–500.

15. Baguelin M, Flasche S, Camacho A, Demiris N, Miller E, Edmunds WJ. Assessing optimal target populations for influenza vaccination programmes: An evidence synthesis and modelling study. PLOS Medicine 2013; 10(10):e1001 527, doi:10.1371/journal.pmed.1001527.

16. Meyer S, Held L. Incorporating social contact data in spatio-temporal models for infectious disease spread. Biostatistics 2016; doi:10.1093/biostatistics/kxw051. Advance Access.

17. Gneiting T, Katzfuss M. Probabilistic forecasting. Annual Review of Statistics and Its Application 2014; 1(1):125–151, doi:10.1146/annurev-statistics-062713-085831.

18. Moran KR, Fairchild G, Generous N, Hickmann K, Osthus D, Priedhorsky R, Hyman J, Del Valle SY. Epidemic forecasting is messier than weather forecasting: The role of human behavior and internet data streams in epidemic forecast. Journal of Infectious Diseases 2016; 214(suppl 4):S404–S408, doi:10.1093/infdis/jiw375.

19. Gneiting T, Raftery AE. Strictly proper scoring rules, prediction, and estimation. Journal of the American Statistical Association 2007; 102(477):359–378, doi:10.1198/016214506000001437.

20. Czado C, Gneiting T, Held L. Predictive model assessment for count data. Biometrics 2009; 65(4):1254–1261, doi:10.1111/j.1541-0420.2009.01191.x.

21. Seillier-Moiseiwitsch F, Sweeting TJ, Dawid AP. Prequential tests of model fit. Scandinavian Journal of Statistics 1992; 19(1):45–60.

22. Seillier-Moiseiwitsch F, Dawid AP. On testing the validity of sequential probability forecasts. Journal of the American Statistical Association 1993; 88(421):355–359, doi:10.1080/01621459.1993.10594328.

23. Wei W, Held L. Calibration tests for count data. Test 2014; 23(4):787–805, doi:10.1007/s11749-014-0380-8.

24. Gneiting T, Stanberry LI, Grimit EP, Held L, Johnson NA. Assessing probabilistic forecasts of multivariate quantities, with an application to ensemble predictions of surface winds. Test 2008; 17(2):211–235, doi:10.1007/s11749-008-0114-x.

25. Scheuerer M, Hamill TM. Variogram-based proper scoring rules for probabilistic forecasts of multivariate quantities. Monthly Weather Review 2015; 143(4):1321–1334, doi:10.1175/MWR-D-14-00269.1.

26. Wei W, Balabdaoui F, Held L. Calibration tests for multivariate Gaussian forecasts. Journal of Multivariate Analysis 2017; 154:216–233, doi: 10.1016/j.jmva.2016.11.005.

27. Held L, Höhle M, Hofmann M. A statistical framework for the analysis of multivariate infectious disease surveillance counts. Statistical Modelling 2005; 5(3):187–199, doi:10.1191/1471082X05st098oa.

28. Meyer S, Held L, Höhle M. Spatio-temporal analysis of epidemic phenomena using the R package surveillance. Journal of Statistical Software ; URL http://arxiv.org/abs/1411.0416, to appear.

29. Pringle K, Lopman B, Vega E, Vinje J, Parashar UD, Hall AJ. Noroviruses: Epidemiology, immunity and prospects for prevention. Future Microbiology 2015; 10(1):53–67, doi:10.2217/fmb.14.102.

30. Paul M, Held L, Toschke A. Multivariate modelling of infectious disease surveillance data. Statistics in Medicine 2008; 27(29):6250–6267, doi: 10.1002/sim.3440.

31. Held L, Paul M. Modeling seasonality in space-time infectious disease surveillance data. Biometrical Journal 2012; 54(6):824–843, doi:10.1002/bimj.201200037.

32. Brockmann D, Hufnagel L, Geisel T. The scaling laws of human travel. Nature 2006; 439(7075):462–465, doi:10.1038/nature04292.

33. Cliff AD, Ord JK. Model building and the analysis of spatial pattern in human geography. Journal of the Royal Statistical Society. Series B (Methodological) 1975; 37(3):297–348.

34. Mossong J, Hens N, Jit M, Beutels P, Auranen K, Mikolajczyk R, Massari M, Salmaso S, Tomba GS, Wallinga J, et al.. Social contacts and mixing patterns relevant to the spread of infectious diseases. PLoS Medicine 2008; 5(3):e74, doi:10.1371/journal.pmed.0050074.

35. Wallinga J, Teunis P, Kretzschmar M. Using data on social contacts to estimate age-specific transmission parameters for respiratory-spread infectious agents. American Journal of Epidemiology 2006; 164(10):936–944, doi:10.1093/aje/kwj317.

36. Bland JM, Altman DG. Statistical methods for assessing agreement between two methods of clinical measurement. The Lancet 1986; 327(8476):307–310, doi:10.1016/S0140-6736(86)90837-8.

37. Good IJ. Rational decisions. Journal of the Royal Statistical Society. Series B (Methodological) 1952; 14(1):107–114.

38. Dawid AP, Sebastiani P. Coherent dispersion criteria for optimal experimental design. Annals of Statistics 1999; 27(1):65–81.

39. Riebler A, Held L. Projecting the future burden of cancer; Bayesian age-period-cohort analysis ready for routine use. Biometrical Journal ; Accepted.

40. Diebold FX, Gunther TA, Tay AS. Evaluating density forecasts with applications to financial risk management. International Economic Review 1998; 39(4):863–883, doi:10.2307/2527342.

41. Diebold FX, Mariano RS. Comparing predictive accuracy. Journal of Business & Economic Statistics 1995; 13(3):253–263, doi:10.1080/07350015.1995.10524599.

42. Hemri S, Lisniak D, Klein B. Multivariate postprocessing techniques for probabilistic hydrological forecasting. Water Resources Research 2015; 51(9):7436–7451, doi:10.1002/2014WR016473.

43. Claeskens G, Hjort NL. Model Selection and Model Averaging. Cambridge University Press: Cambridge, 2008.

44. Sloughter M, Gneiting T, Raftery AE. Probabilistic wind speed forecasting using ensembles and Bayesian model averaging. Journal of the American Statistical Association 2010; 105(489):25–35, doi:10.1198/jasa.2009.ap08615.

